# Aerosphere occupancy (ψ) model of lesser short nosed fruit bat (*Cynopterus brachyotis* Muller, 1838) related to tree species in Asia mountainous paddy fields

**DOI:** 10.1101/2021.02.08.430234

**Authors:** Adi Basukriadi, Erwin Nurdin, Andri Wibowo, Jimi Gunawan

## Abstract

Bat is animal that occupies aerosphere, especially fruit bats that forage on the space around the trees. The fruit bats use whether narrow space below tree canopy or in edge space on the edge of canopy. Whereas the aerosphere occupancy of fruits bats related to the specific tree species is poorly understood. Here, this paper aims to assess and model the association of fruit bat *Cynopterus brachyotis* aerosphere occupancy (Ψ) with tree species planted in mountainous paddy fields in West Java. The studied tree species including *Alianthus altissima, Acacia* sp., *Cocos nucifera, Mangifera indica, Pinus* sp., and *Swietenia macrophylla*. The result shows that the tree species diversity has significantly (x^2^= 27.67, P < 0.05) affected the *C. brachyotis* aerosphere occupancy. According to values of Ψ and occupancy percentage, high occupancy of narrow space by *C. brachyotis* was observed in *Swietenia macrophylla* (Ψ = 0.934, 78%), followed by *Alianthus altissima* (Ψ = 0.803, 57%), and *Mangifera indica* (Ψ = 913, 55%). While high occupancy of edge space was observed in *Mangifera indica* (Ψ = 0.685, 41%), followed by *Pinus* sp. (Ψ = 0.674, 38%), and *Alianthus altissima* sp. (Ψ = 0.627, 36%). The best model for explaining *C. brachyotis* occupation in narrow space is the tree height with preferences on high tree (Ψ~tree height, AIC = 1.574, R^2^= 0.5535, Adj. R = 0.4047). While for edge space occupant, the best model is also the tree height (Ψ~tree height, AIC = −26.1510, R^2^= 0.7944, Adj. R = 0.7258).

## INTRODUCTION

Bat assemblages are influenced by the differential uses of the aerosphere that can be classified into 3 spaces (Figure 1) and habitat types including open space that far away from obstacles or the ground, edge space close to, but not within, vegetation or above water, and narrow space within vegetation and under the canopy layer. Each of these space partitions of the aerosphere related to particular adaptations in shape of the wings and in design of echolocation calls. Bats flying and using open space are characterized by long and narrow wings (Marinello and Bernard 2014, Norberg and Rayner 1987) that permit fast flight. This bat species usually has low frequency and narrowband echolocation calls that are an adaptation for long-range detection of insects. Bats hunting closer to obstacles, foliage, and using narrow space are characterized by broader and somewhat shorter wings. Echolocation calls have higher frequency. Wing shapes of bats that mostly fly within the forest and use narow space under tree canopy are broad and short to allow skilful maneuvering around obstacles and tree branches (Schnitzler and Kalko 2001, Korine and Kalko 2005).

**Figure 1.**
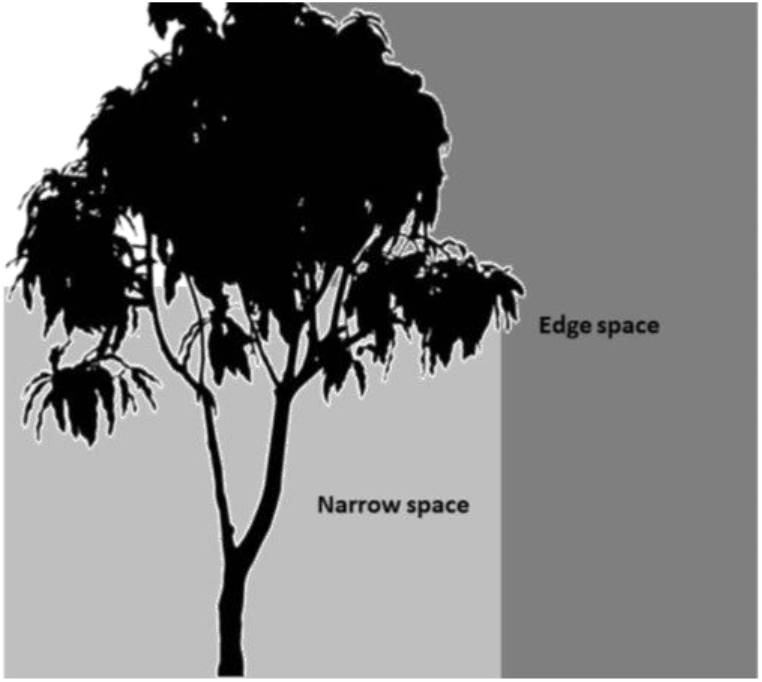
Space use of bats, narrow space is space under tree canopy and edge space is space on the edge of tree canopy.

Bat and tree species preferences are widely studied whereas the exact space occupancy of bats related to the particular tree species is still poorly understood. Asia is one of region that has high diverse of tree species that is important to support the arboreal bats including fruit bat *C. brachyotis*. Here this paper aims to model the *C. brachyotis* occupancy on space surrounding trees.

## MATERIALS AND METHODS

### Study area

This study was conducted in mountainous landscape located in Nagreg region in West Java Province, Indonesia. This landscape was dominated by mix of paddy fields, plantations, and settlements (Figure 2). The elevation was between 718 to 728 m above sea levels. In this study area, 3 transects with length of each transect was 1 km were located. The fruit bat, vegetation structure and aerosphere variable surveys were conducted in those transects in August 2020.

**Figure 2.**
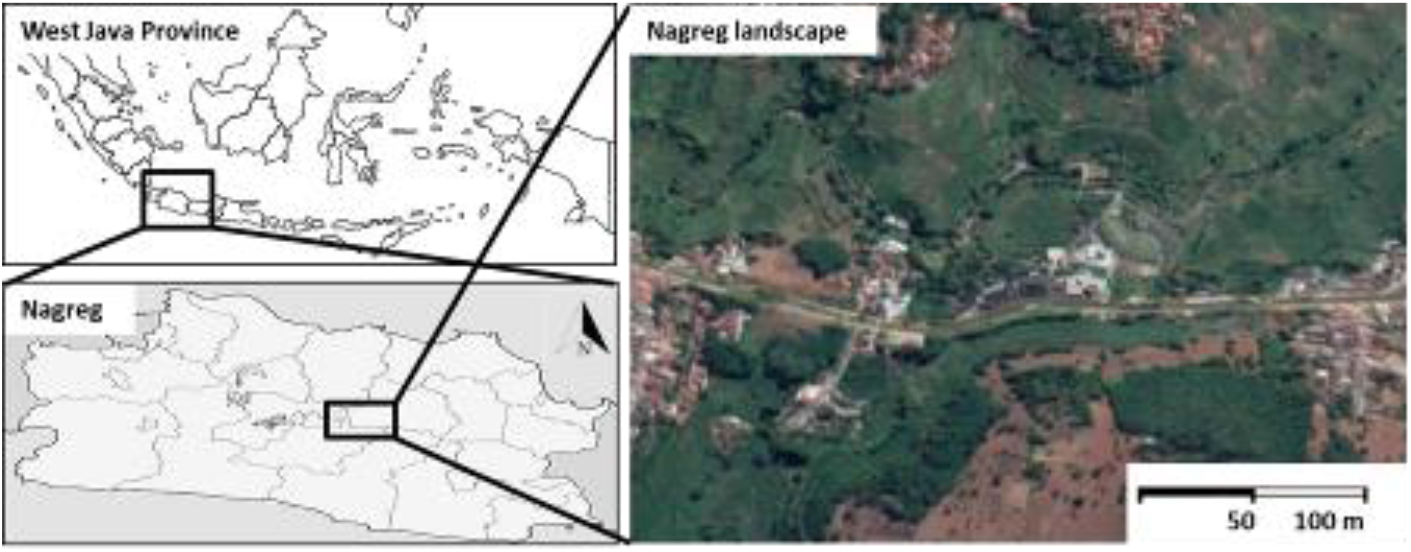
Location of study area in Nagreg landscape in West Java Province, Indonesia.

### Method

#### Tree species assemblage surveys

Tree surveys were conducted in August 2020 following the fruit bat surveys from 17.00 to 19.00. The trees that were passed and perched by bats were recorded. The recorded variables were including tree species, tree canopy cover and height. The surveys were conducted within a square plot sizing 10 m x 10 m standardized for tree surveys.

#### Cynopterus brachyotis surveys

*C. brachyotis* observations were conducted following methods as described by several authors. The observations were conducted at noon from 17.00 to 19.00 following *C. brachyotis* activity times within 10 m × 10 m grid in each sampling location. *C. brachyotis* were recorded and denoted as presence and absence data with 3 sampling event replications. *C. brachyotis* observations were emphasizing on bat presences in space below the tree canopy (narrow space) and space on the edge of tree canopy (edge space).

### Data analysis

Aerosphere occupancy analysis was following Coleman et al. (2014) and Starbuck et al. (2015). The occupancy analysis was performed upon comparisons of events with presences of bats and total sampling events. The occupancy variables were denoted as occupancy (Ψ), occupancy percentage (%), detection probability (*p*) and naïve estimates. The occupancy analyses were performed to compare bat presence in narrow and edge spaces as functions of tree species. The significance difference of bat occupancy affected by tree species was analyzed using x^2^ test with significance level at P < 0.05.

Occupancy model as functions of tree cover and height was developed using Akaike Information Criterion (AIC). The AIC was developed using the linear regression. The measured parameters included in AIC are R^2^ and adjusted R^2^. To build the model, 2 explanatory covariates including tree covers in %/100 m^2^, tree heights and combinations of those covariates were included in the analysis to develop the model.

## RESULTS AND DISCUSSION

### Space occupancy selection

There were 6 tree species preferred by *C. brachyotis* including *Alianthus altissima, Acacia* sp., *Cocos nucifera, Mangifera indica, Pinus* sp., and *Swietenia macrophylla*. In those tree species, *C. brachyotis* was observed foraging in edge space and narrow space. The results show that *C. brachyotis* use narrow space more frequently than edge space (Figure 3). The space uses were affected (x^2^= 27.67, P < 0.05) by tree species. According to values of Ψ and occupancy percentage, high occupancy of narrow space by *C. brachyotis* was observed in *Swietenia macrophylla* (Ψ = 0.934, 78%), followed by *Alianthus altissima* (Ψ = 0.803, 57%), and *Mangifera indica* (Ψ = 913, 55%). While high occupancy of edge space was observed in *Mangifera indica* (Ψ = 0.685, 41%), followed by *Pinus* sp. (Ψ = 0.674, 38%), and *Alianthus altissima* sp. (Ψ = 0.627, 36%) (Table 1). The best model for explaining *C. brachyotis* occupation in narrow space is the tree height with preferences on high tree (Ψ ~tree height, AIC = 1.574, R^2^ = 0.5535, Adj. R = 0.4047). While for edge space occupant, the best model is also the tree height (Ψ ~tree height, AIC = −26.1510, R^2^= 0.7944, Adj. R = 0.7258) (Table 2).

**Figure 3.**
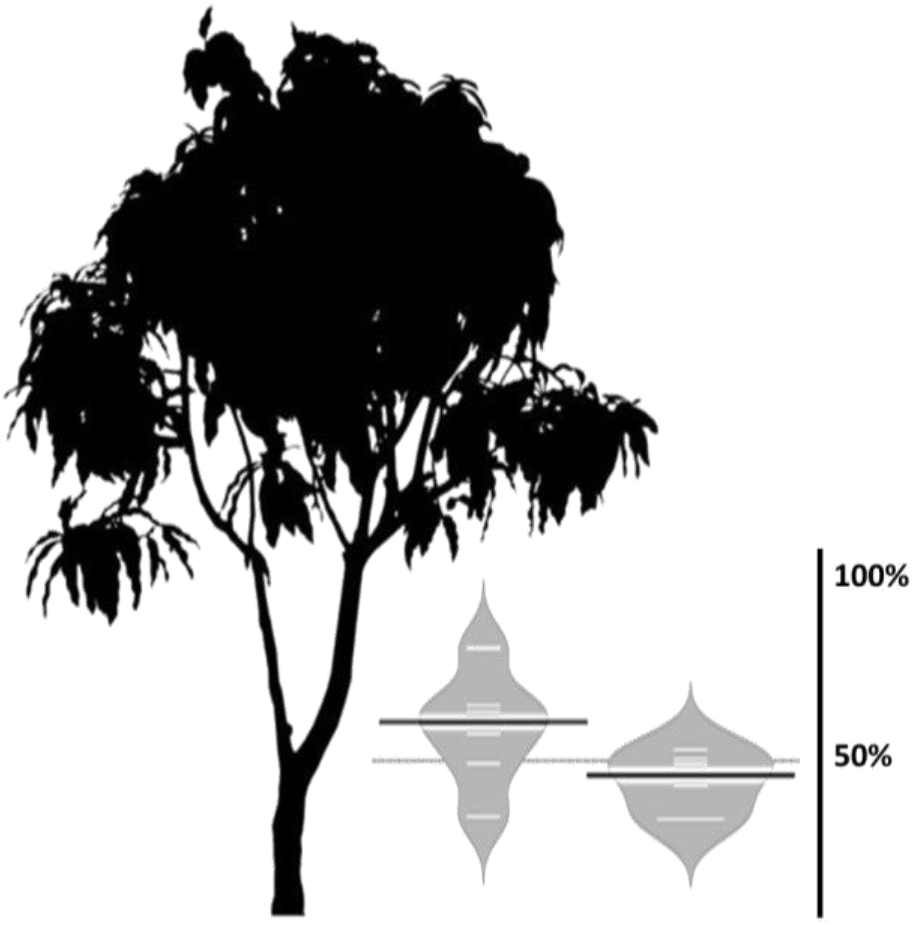
Space occupancy percentage of *C. brachyotis* and comparison between narrow and edge spaces (x^2^= 27.67, P < 0.05)

**Table 1.**
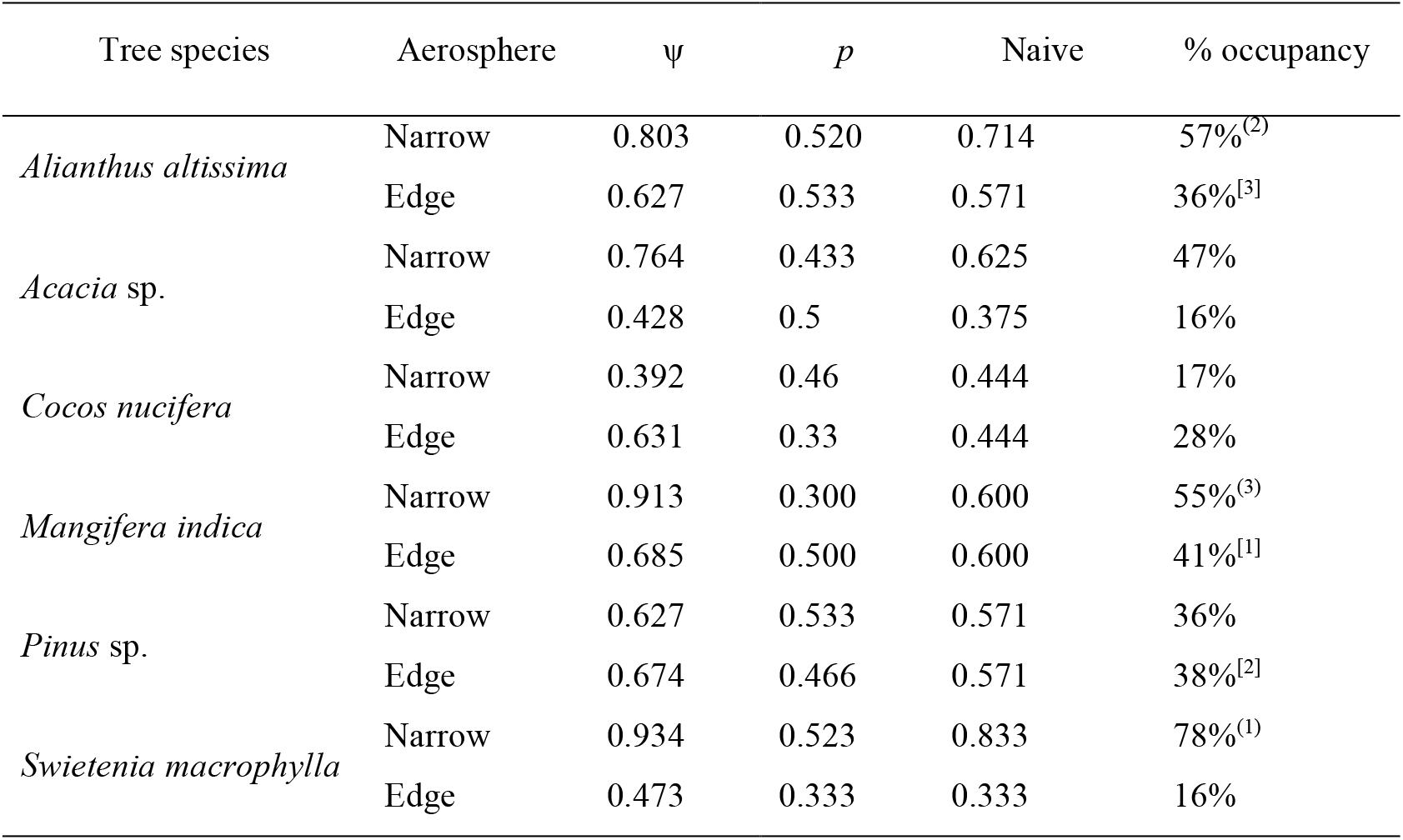
Values of habitat occupancy (Ψ), detection probability (*p*) and occupancy percentage of fruit bat *Cynopterus brachyotis* in *Alianthus altissima, Acaci*a sp., *Cocos nucifera, Mangifera indica, Pinus* and *Swietenia macrophylla* trees. Superscript numbers in bracket show order of values from high to low.

| Tree species | Aerosphere | $\psi$ | $p$ | Naive | % occupancy |
| --- | --- | --- | --- | --- | --- |
| *Alianthus altissima* | Narrow | 0.803 | 0.520 | 0.714 | 57%<sup>(2)</sup> |
|  | Edge | 0.627 | 0.533 | 0.571 | 36%<sup>[3]</sup> |
| *Acacia* sp. | Narrow | 0.764 | 0.433 | 0.625 | 47% |
|  | Edge | 0.428 | 0.5 | 0.375 | 16% |
| *Cocos nucifera* | Narrow | 0.392 | 0.46 | 0.444 | 17% |
|  | Edge | 0.631 | 0.33 | 0.444 | 28% |
| *Mangifera indica* | Narrow | 0.913 | 0.300 | 0.600 | 55%<sup>(3)</sup> |
|  | Edge | 0.685 | 0.500 | 0.600 | 41%<sup>[1]</sup> |
| *Pinus* sp. | Narrow | 0.627 | 0.533 | 0.571 | 36% |
|  | Edge | 0.674 | 0.466 | 0.571 | 38%<sup>[2]</sup> |
| *Swietenia macrophylla* | Narrow | 0.934 | 0.523 | 0.833 | 78%<sup>(1)</sup> |
|  | Edge | 0.473 | 0.333 | 0.333 | 16% |

**Table 2.**
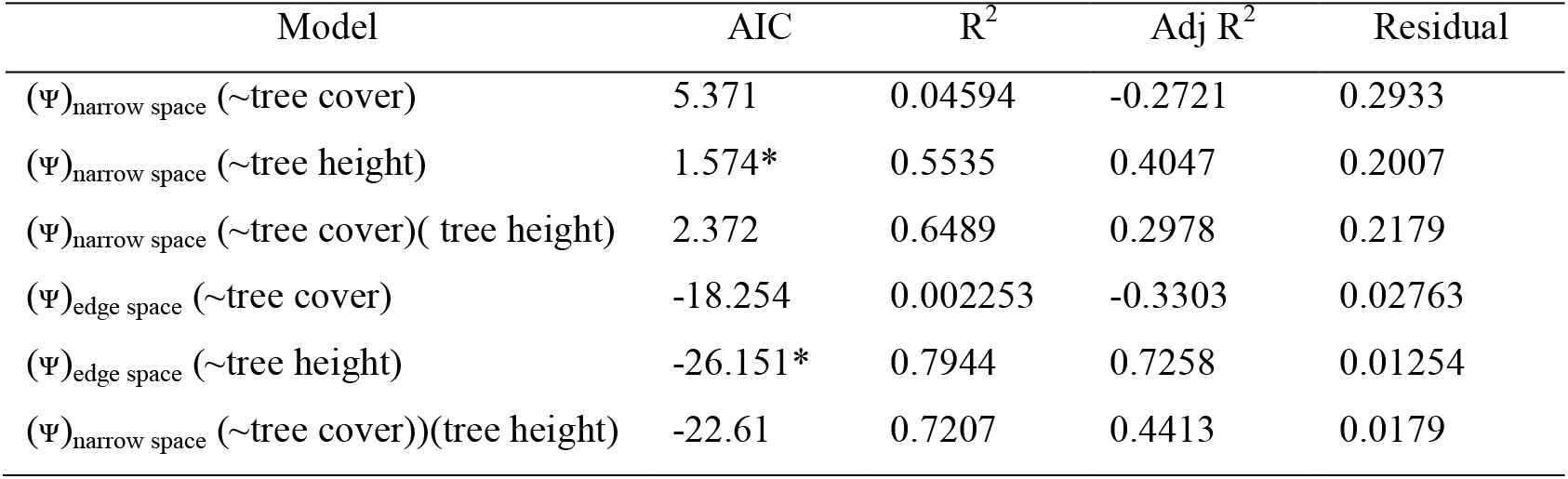
Selected occupancy model (asterisk signs) of fruit bat *Cynopterus brachyotis* with tree cover-height covariates.

### Discussion

The results show significant occupancy of fruit bat on narrow space below canopy. This space was known preferred by bat species. Kalko and Handley (2001) observed that canopy has more bat species with the canopy rigs yielded more species (n = 41) than the ground nets (n = 35). Bats known use the canopy for a variety of purposes including roosting, foraging, and reproduction. The use of canopy was 76% for roosting and 85% for foraging (Wunder and Carey 1996). A space below canopy tree provides several features that can support the bats. Those features are including tree cavities and foliage. Tree cavities used by bats including hollows formed in the trunks or branches of snags or damaged live trees. Tree cavities generally provide a relatively stable microclimate and offer protection from predators. Several factors influencing bat occupancies in space below canopy are microclimate, structure, tree age, size, and height. The most important features of narrow space are the foliage. Narrow space with its foliage above is generally well insulated against cold temperatures and reducing predator risks.

Whereas in our study, the *C. brachyotis* shows opposite patterns of space occupancy in one particular trees species. In contrast to other tree species, the space occupancy for *Cocos nucifera* was higher in edge rather than in narrow space. While in general, space occupancy of *C. brachyotis* were always higher in narrow space rather than in edge space. This condition was related to the morphology of *C. nucifera* tree in particular its canopy. This species is not belong to the hard wood species even it has height almost equal to other tree species. Since it is not belong to hard wood species, this species does not have canopy cover as dense as canopy cover belongs to hard wood species. Beside effects related to canopy covers, some bat species were observed preferring more edge space above canopy rather than narrow space below canopy. Menzel et al. (2005) reported that the mean of bat activities can be 3 folds greater in above canopy in comparison to activity in below canopy.

Bats in Nagreg landscape have high occupancy in mango (*Mangifera indica*) trees. The preferences of bats with mango trees observed in this study were in corroborated with results from other studies. Tollington et al. (2019) reported that the consumption of 31000 lychee fruits were 42% consumed by the Mauritius fruit bat (*Pteropus niger*). From 81 fruit tree species, mango is frequently consumed by bats (Aziz et al. 2016). In West Java regions, Suyanto (2002) reported that occupancy of *Cynopterus* spp. to certain tree species related to the foraging for leaves and fruits as these were main food sources for fruit bats.

In Nagreg, fruit bat also has high occupancy on other non-fruiting tree species other than mango. Soegiharto et al. (2010) have reported several vegetation species that may attract fruit bat presences. Those species were including families of Acanthaceae, Apocynaceae, Begoniaceae, Convolvulaceae, Paceae, Pinaceae, and Orchidaceae, The preferred species were *Anacardium* sp., *Adenanthera* sp., *Syzygium s*p., *Anacardium* sp., *Annona* sp., *Ceiba* sp., *Eugenia* sp., *Hisbiscus* sp., *Acacia* sp., *Cyathea* sp., *Salacia* sp., *Cyperus* sp., and *Croton* sp. In Nagreg, *Acacia* sp. was one of species occupied by *C. brachyotis* with low frequency. In the model, the tree height covariate was the best estimate for bat space occupancy. The converse correlation of height with bat activity was also reported in other studies. Müller et al. (2013) have observed that bat activity was correlated with height of tree.

Tree stands are important for bat population mainly fruit bats. Whereas rationale for selecting appropriate tree species is still challenging. Here this study has employed habitat occupancy method and model to assess tree species used mostly by fruit bats. To conclude, several tree species including *Alianthus altissima*, *Mangifera indica*, and *Swietenia macrophylla* are occupied more frequently by *C. brachyotis* including in their space below canopy and in their space on the edge of canopy.

## Abbreviations

AIC: (Akaike Information Criterion)

